# Conditioned expectations drive false odor detection

**DOI:** 10.64898/2026.11.20.768152

**Authors:** Lauren A. Wolters, Xiaolin Qiao, Laura Shanahan, James D. Howard

**Author notes:** **Corresponding author(s):** James D. Howard.

## Abstract

Research in auditory and visual fields has shown that false percepts can emerge when perceptual priors outweigh incoming sensory information. However, the role of similar top-down mechanisms in generating false percepts in the olfactory domain is less understood. To address this, we implemented a Pavlovian conditioning paradigm in which we first individually determined the odor detection thresholds of Butanol (*n* = 55) using an adaptive algorithm. Participants then completed an odor detection task where on each trial one of two distinct visual cues was paired with either a supra-threshold, threshold, or sub-threshold concentration of Butanol, or no odor. In the first three trial blocks, one visual stimulus (“strong cue”) was paired more frequently with supra-threshold concentrations, and the other visual stimulus (“weak cue”) was paired more frequently with sub-threshold concentrations. In the last three blocks both cues were paired most frequently with the “no odor” condition. We observed significantly greater odor detection rates on “no odor” trials for the strong cue compared to the weak cue. Importantly, this finding could not be explained by differences in total number of visual cue presentations or other task variables. Further, the difference in odor detection rate for the strong and weak cues during the supra-threshold conditioning trials predicted the difference in false odor detection during the “no odor” trials, suggesting that the prior formed between the strong cue and odor contributed to the false olfactory percepts. This demonstrates that false odor perception can be enhanced by the expectation of an olfactory stimulus in a Pavlovian conditioning paradigm. These findings could yield therapeutic insight into the mechanisms underlying pathological olfactory hallucinations.

## Introduction

Perception is increasingly recognized to be an inferential process in which sensory information is influenced by prior experiences and expectations (Friston, 2010). Constructing an internal model that can flexibly update learned associations based on incoming sensory stimuli allows for efficient navigation of complex sensory environments (Kok et al., 2012; Knill and Pouget, 2004; Shaw et al., 2026). Within the predictive coding framework, perception relies on the integration of internal predictions, or priors, and sensory input (Rao & Ballard, 1999; Chen et al., 2025). Discrepancies between the two, known as prediction errors, serve to refine and improve this internal model (Sterzer et al., 2018). However, if a prior outweighs incoming sensory evidence, particularly when the sensory input is ambiguous or below perceptual threshold, the strong prior can dominate resulting in false perceptual experiences (Haarsma et al., 2023; Koh & Gallagher, 2020).

Research in the auditory and visual domains have demonstrated that false percepts can arise when perceptual priors are overweighted relative to incoming sensory evidence (Kafadar et al., 2022; Kok et al., 2012; Nazimek et al., 2012; Powers et al., 2017; Shaw et al., 2026). However, whether similar mechanisms produce false percepts in olfaction remains largely unexplored despite growing evidence that olfactory perception is highly sensitive to contextual and expectation-driven modulation (Lyons & Gottfried, 2025; Zelano et al., 2011).

The olfactory system is unique among sensory modalities due to its strong links with memory and emotion, involving complex interactions between sensory input and high-order brain areas such as the orbitofrontal cortex and piriform cortex (Gottfried & Dolan, 2003; Plailly et al., 2008). While studies have demonstrated that verbal labels, visual context, and associative learning can alter odor perception, affecting intensity, discriminability and hedonic valence (de Araujo et al., 2005; Djordjevic et al., 2008; Li et al., 2008), the question of whether top-down effects can induce the perception of odors in the absence of physical stimuli remains largely unanswered. Investigating this phenomenon is crucial not only for advancing our understanding of the neural mechanisms of olfaction but also for its relevance to clinical conditions characterized by maladaptive olfactory distortions or hallucinations.

To explore how false percepts arise among individuals, we developed an odor detection task investigating top-down influences on odor perception. Drawing from the well-established Conditioned Hallucination Paradigm (Powers et al., 2017), our task was designed to reliably build an association between visual stimuli and an odor, Butanol. Notably however, we employed two distinct conditioned stimuli, one of which was paired more frequently with the odor during the initial phase of the experiment. This asymmetrical pairing was designed to establish a strong associative prior for one visual stimulus relative to the other. In the later stages of the task, both stimuli were primarily presented without any odor. This allowed us to test whether the learned association alone was sufficient to elicit false odor percepts under controlled experimental conditions. This novel approach directly examines whether top-down expectations contribute to false sensory experiences in the olfactory domain.

Based on prior work, we hypothesized that the association between a strongly conditioned visual stimulus and an odor would lead individuals to report smelling an odor when none was delivered. Further we aimed to explore whether this effect was mediated by a higher predisposition to olfactory mental imagery in daily life or delusional tendencies.

## Materials & Methods

### Participants

At total 55 participants (18 male, 34 female, 3 other) were recruited from the student population at Brandeis University to take part in this study. All participants were fluent English speakers between the ages of 18 and 39 (*M*_age_ ± *SD* = 21.13 ± 4.63 years). Participants provided informed consent and self-reported to be right-handed non-smokers with no history of significant neuropsychiatric disorder, epilepsy, or migraine/headache disorder. Participants reported to not be currently taking psychotropic medication, and had no known visual or olfactory impairments. Participants were compensated $15 or one course credit according to the Brandeis University Research Familiarization process. The study protocol was approved by the Brandeis University Institutional Review Board.

### Visual Stimuli

Twelve distinct abstract fractal images were selected from a standardized database to be used in this study (Ovalle-Fresa et al., 2021). For each participant, two fractal images were randomly chosen from the set of twelve to be used as conditioned cues for the duration of the experiment.

### Odor Stimuli

The olfactory stimulus, Butanol, is a commercially available odorant produced by the Flavors and Fragrances Division of Sigma-Aldrich Fine Chemicals. Odorants in this category are intended for use as flavor or perfume substances and therefore meet specifications in the Food Chemicals Codex (Fifth Edition, 2003) for human exposure and consumption. Butanol is commonly used in standardized olfactory psychophysical tests, such as the widely used ‘Sniffin’ sticks’ (Hummel et al., 1997). The odorant was diluted with mineral oil to produce a range of eight concentrations for threshold testing. For 5 of the 55 participants reported here, the set of concentrations used ranged from 0.0625% (strongest) to 0.00000024% (weakest) in dilution steps of ¼. For 50 of the 55 participants the set of concentrations used ranged from 0.25% (strongest) to 0.00000012% (weakest) in dilution steps of ⅛.

### Odor presentation

The odorant was delivered using a custom-built olfactometer that directs medical-grade room air via Teflon tubes and mass flow controllers (Alicat Scientific, Tucson, AZ) through the headspace of glass bottles containing approximately 3.5 mL of liquid odorant. Using custom MATLAB (MathWorks, Natick, MA) scripts, the olfactometer was programmed to switch between un-odorized and odorized air with precise timing and flow rates through any individual bottle or combination of bottles. In this system, air converges on a single Teflon tube that terminates in a nasal mask (Philips Respironics, Murrysville, PA) worn by the participant. To avoid somatosensory stimulation in the nose, a constant airflow rate of 3 L/min was maintained throughout the experiment, and odorized air was mixed into the un-odorized air stream without a change in total flow.

### Physiological Measurements

Respiration was measured throughout the experiment using a spirometer (ADInstruments, Colorado Springs, CO) connected to the nasal mask. This device measures nasal inspiration/expiration with a flow sensor connected to a PowerLab 8 Channel recorder (ADInstruments) which converts the signal to a digital signal for later analysis via LabChart software.

### Experiment Design

This experiment consisted of a brief online screening and questionnaire session, followed by a 50-minute in-person testing session conducted at a later date in a behavioral testing room at Brandeis University.

### Questionnaires

At the beginning of the experiment participants completed four standardized questionnaires designed to assess anxiety and depression (Hospital Anxiety and Depression Scale, HADS; Zigmond & Snaith, 1983), experience of hallucinations (Launay-Slade Hallucination Scale – Extended, LSHS-E; Larøi & Van Der Linden, 2005), delusions (Peters et al. Delusions Inventory, PDI; Peters et al., 2004), and olfactory mental imagery (Vividness of Olfactory Imagery Questionnaire, VOIQ; Gilbert et al., 1998).

### Ratings

Participants next provided ratings of pleasantness (anchors “extremely unpleasant” and “extremely pleasant”) and familiarity (anchors “extremely unfamiliar” and “extremely familiar”) on a visual analog scale for the strongest concentration of Butanol used in the odor dilution series. At the end of the experiment, participants were asked to rate the likelihood that each visual cue was paired with the odor on a visual analog scale from 1 (sure no odor) to 5 (sure odor).

### Threshold Task

The threshold task consisted of 24 total trials. On each trial one visual cue was presented simultaneously with one of the odor concentrations in the dilution series. The concentration of odor delivered on each trial was determined by an adaptive QUEST algorithm (Watson & Pelli, 1983), implemented via Psychtoolbox in MATLAB. Participants were instructed to make a Yes/No response using the left and right arrow keys on a standard keyboard as to whether or not they detected the odor on each trial. Twenty trials have shown to reliably identify an odor threshold using an adaptive psychometric algorithm in prior studies (Höchenberger & Ohla, 2019), and four additional trials were used here to ensure that every odor concentration was presented at least once during the threshold task.

A standard Weibull psychometric function was fitted to individual participant responses to determine the odor concentration at which 50% of Yes responses were made, rounded to the nearest discrete number in our odor set. The supra-threshold concentration was then set as 2 steps stronger than threshold, and the sub-threshold concentration was set as 2 steps weaker than threshold. If the calculated threshold was less than our lowest possible concentration step (i.e. concentration step 3) or greater than our highest possible concentration step (i.e. concentration step 6), we forced the threshold to be step 3 or step 6, respectively. Participants were not excluded from the data set based on their determined threshold concentration.

### Odor Detection Task

The subsequent odor detection task proceeded in 6 blocks of 20 trials each. On each trial one of the two visual cues was presented simultaneously with delivery of either the supra-threshold, threshold, or sub-threshold concentration of the odor (as determined by the threshold task), or no odor. Participants were instructed to sniff when they saw the image and then make a Yes/No response as to whether they detected an odor. Additionally, they were instructed to hold the response key down to indicate, with visual feedback on a sliding scale, how confident they felt about their response. Longer key press times corresponded to higher confidence, indicated by higher ratings on the sliding scale.

Participants were not made explicitly aware of differences in the likelihood of receiving a supra-threshold odor in conjunction with either visual cue. Covertly, we designated one of the two cues as the “strong” conditioned stimulus (CSs, see **Fig. 2a**), as it was more frequently paired with the supra-threshold odor concentration in the first three blocks of the task. The other cue was designated as the “weak” conditioned stimulus (CSw), as it was more frequently paired with threshold and sub-threshold concentrations in the first three blocks. Both cues were paired with the same ratio of odor concentrations in the last three blocks of the task. Because there were more CSs trials than CSw trials in the first three blocks, we included the CSw in all trials of the threshold task to ensure that the total number of presentations of CSs and CSw was equal (72 times each) across the entire experiment.

### Respiration Analysis

Respiration was analyzed in MATLAB using custom scripts. We tested for potential differences in cued sniffing in the “no odor” condition for the CSs and CSw. For each trial, the sniff peak was calculated as the maximum signal amplitude in the 3s after the sniff cue minus the signal amplitude at the time of sniff cue. Sniff duration was calculated as the time from sniff cue to sniff peak. Peak and duration were averaged across trials and submitted to paired *t*-tests for hypothesis testing.

### miniPID Analysis

To verify that we achieved separable odor concentrations, and no odor delivery on putative no-odor trials, we conducted an independent test of olfactometer performance using a photoionization detector (PID; Aurora Scientific, Ontario, Canada). The sensor head was placed at the end of the single Teflon tube and subjected to the same 120 total instances of odor or no-odor delivery: 24 each of the subject-average supra-threshold, threshold, and sub-threshold concentration steps, and 48 instances of no odor delivery. Sensory responses were averaged across instances and cumulative responses were submitted to ANOVA and *t*-tests for hypothesis testing.

### Statistical Analysis

Statistical analyses were performed in MATLAB. We used logistic regression to model binary odor detection responses as a function of current and previous trial variables. Paired-samples *t*-tests were used to compare odor detection rates, confidence ratings, response times, sniff parameters, and odor likelihood ratings across cue conditions and response types. Pearson correlation coefficients were calculated to examine relationships between conditioning effects and false odor detection rates, and Spearman rank correlation was used to assess the relationship between odor likelihood ratings and false odor detection rate. Correlation analyses were also used to examine associations between false odor detection and questionnaire measures. Statistical significance was set at *p* < 0.05. Trials in which no response was made were excluded from analysis. All 55 included participants passed a quality-control criterion of fewer than 10% missed responses during the odor detection task.

## Results

### Stimulus selection and odor threshold determination

For each participant, two fractal images were randomly chosen from a set of 12 to be used as conditioned cues in both the initial odor threshold task and subsequent odor detection task (**Fig. 1A**). The odor stimuli used in these tasks consisted of an 8-step serial dilution of Butanol in mineral oil (**Fig. 1B**). On each trial of the odor threshold task, one of the two visual cues was paired with one of the 8 Butanol concentrations and participants were instructed to make a Yes/No odor detection response. (**Fig. 1C**).

**Figure 1.**
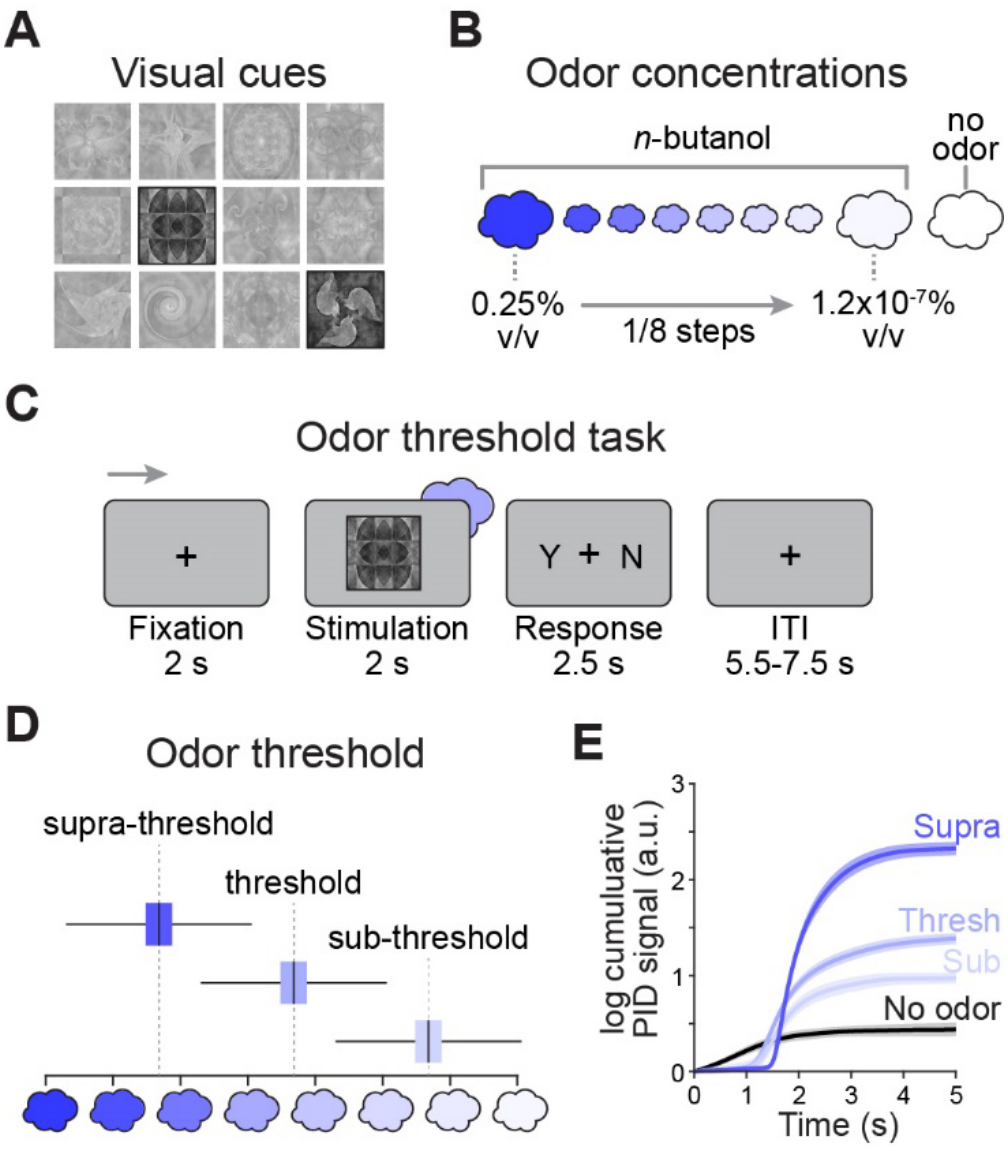
Stimulus selection and odor threshold testing. **(A)** Example of two fractal images randomly selected for one participant from the set of 12 to serve as visual conditioned stimuli for the remainder of the experiment. **(B)** Butanol was dissolved in odorless mineral oil in 1/8 steps to create an 8-step dilution series. Darker shades of blue correspond to stronger concentrations. The odor detection task also included a “no odor” condition. **(C)** On each trial of the odor threshold task participants were presented simultaneously with one of the two CS’s and one concentration of the dilution series. Participants were instructed to respond “Yes” if they detected the Butanol odor and “No” if they did not. **(D)** The average odor threshold concentration calculated across participants fell in the middle of the dilution series. Supra-threshold and sub-threshold concentrations were set as 2 steps stronger and 2 steps weaker than threshold, respectively. Error bars depict standard deviation, and the extent of the boxes depict standard error of the mean. **(E)** Cumulative photo-ionization detector responses to 30 presentations each of average supra-threshold, threshold, and sub-threshold concentrations revealed clearly separable response profiles by concentration, and no response on putative “no odor” trials. Our olfactometer is thus capable of consistently delivering intended concentration gradients, even at very low concentration values, without lingering odor on “no odor” trials. Shaded areas surrounding lines depict standard deviation.

Responses from the threshold task were fitted with a Weibull psychometric function, and threshold was determined as the odor concentration step corresponding to 50% probability of a “Yes” response on the fitted curve. The average calculated threshold fell in the middle of the dilution series (*M*_concentration step_ ± *SD* = 4.32 ± 1.38) (**Fig. 1D**). We set the individually determined threshold concentration as this number rounded to the nearest integer, and designated putative “supra-threshold” and “sub-threshold” concentrations as two steps stronger and two steps weaker than the threshold step, respectively (**Fig. 1D**).

We tested whether this procedure resulted in distinct concentrations of odor delivery by conducting an independent analysis of photoionization detector (PID) responses to the average sub-threshold, threshold, and supra-threshold concentration steps. This analysis confirmed clearly separable responses to each concentration step, and no response to the “no odor” condition (**Fig. 1E**).

### Odor detection task design and validation

After completing the odor threshold task, participants next completed 6 blocks of 20 trials each of an odor detection task. In this task one of the cues, denoted here as the “strong” conditioned stimulus (CSs), was paired more frequently with supra-threshold odor concentration in the first 3 blocks of the task than the other cue, denoted here as the “weak” conditioned stimulus (CSw) (**Fig. 2A**). In the last 3 blocks of the task the CSs and CSw were both most frequently paired with the “no odor” condition in equal proportions (**Fig. 2A**). On each trial, after the simultaneous cue and odor presentation participants made a Yes/No odor detection response and held the response key down to indicate in a fillable bar how confident their detection was (**Fig. 2B**). This experimental design allowed us to systematically investigate whether participants made more frequent and more confident false odor detection responses in the CSs condition than the CSw condition.

**Figure 2.**
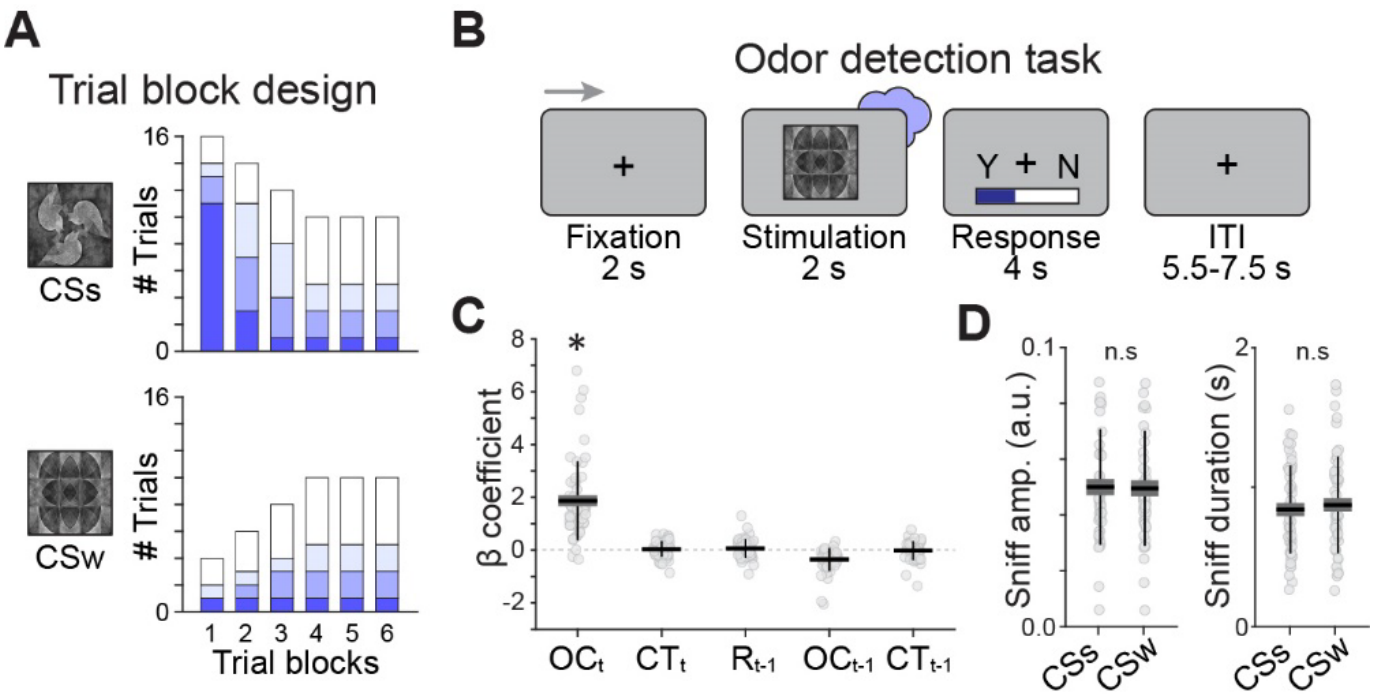
Odor detection task design and validation. **(A)** Histograms for each CS type depict the number of trials in each block paired with the individually calculated supra-threshold (darkest shade), threshold (medium shade), sub-threshold (lightest shade), and no odor (white) concentrations. Each block contained 20 trials. **(B)** The odor detection task was the same as the odor threshold task, except that here participants were instructed to hold the response key down to indicate the confidence of their response (whether “Yes” or “No”). Unfilled bar represents no confidence, completely filled bar represents maximum confidence. **(C)** Regression coefficients are plotted for the five task features included in the logistic regression analysis, modeled on current (t) and previous (t-1) trials with respect to current detection response (OC = odor concentration, CT = cue type, R = response type) **(D)** Sniff amplitude and duration for the CSs vs CSw is plotted for “no odor” trials, which are the main focus of analysis in the odor detection task (n.s. = not significant). In both **(C)** and **(D)**, horizontal lines depict the mean across participants, vertical lines depict standard deviation, the extent of the boxes depict standard error of the mean, and light gray circles depict individual participants.

We first examined which task features generally influenced odor detection responses by conducting a logistic regression analysis with detection responses modeled as a linear function of five features: odor concentration on the current trial (OC_t_); CS type on the current trial (CT_t_), response on the previous trial (R_t-1_), and odor concentration and CS type on the previous trial (OC_t-1_, CT_t-1_). We found a significant position regression coefficient only for the OC_t_ factor (t_54_ = 9.27, p = 9.19×10^−13^), and a significant negative coefficient for the OC_t-1_ factor (t_54_ = 6.06, p = 1.35×10^−7^) (**Fig. 2C**). This analysis confirms that, across all trial types, odor detection responses were primarily influenced by the concentration of the delivered odor, rather than a systematic bias toward one particular cue or response tendency. Moreover, analysis of nasal respiration measured by a spirometer throughout the experiment revealed no significant differences between CSs and CSw on “no odor” trials in sniff amplitude (t_54_ = 0.64, p = 0.52) or sniff duration (t_54_ = −1.33, p = 0.19) (**Fig. 2D**), suggesting that any observed differences in false odor detection rates on these trials were not driven by potential respiratory biases.

### Odor detection task results

We next tested whether participants exhibited different odor detection response rates between CSs and CSw conditions at each of the four odor concentration steps across the entire experiment. There was no difference in odor detection rate between CSs and CSw in the supra-threshold (t_54_ = 0.058, p = 0.95), threshold (t_54_ = 1.39, p = 0.17), or sub-threshold conditions (t_54_ = 0.12, p = 0.91). However, in the “no odor” condition participants reported detecting an odor significantly more frequently on CSs trials than CSw trials (t_54_ = 2.08, p = 0.043) (**Fig. 3A**). Moreover, across participants the difference between CSs and CSw odor detection rate on “no odor” trials was positively correlated with the difference between CSs and CSw odor detection rate on supra-threshold trials (*r* = 0.37, *p* = 0.0052), but not threshold (*r* = −0.0022, *p* = 0.99), or sub-threshold (*r* = −0.011, *p* = 0.93) trials (**Fig. 3B**). Taken together, these findings suggest that the more frequent pairing of the CSs cue with supra-threshold concentrations compared to the CSw cue created a stronger expectation of odor for that condition, resulting in more frequent false odor detections on “no odor” trials for the CSs cue.

**Figure 3.**
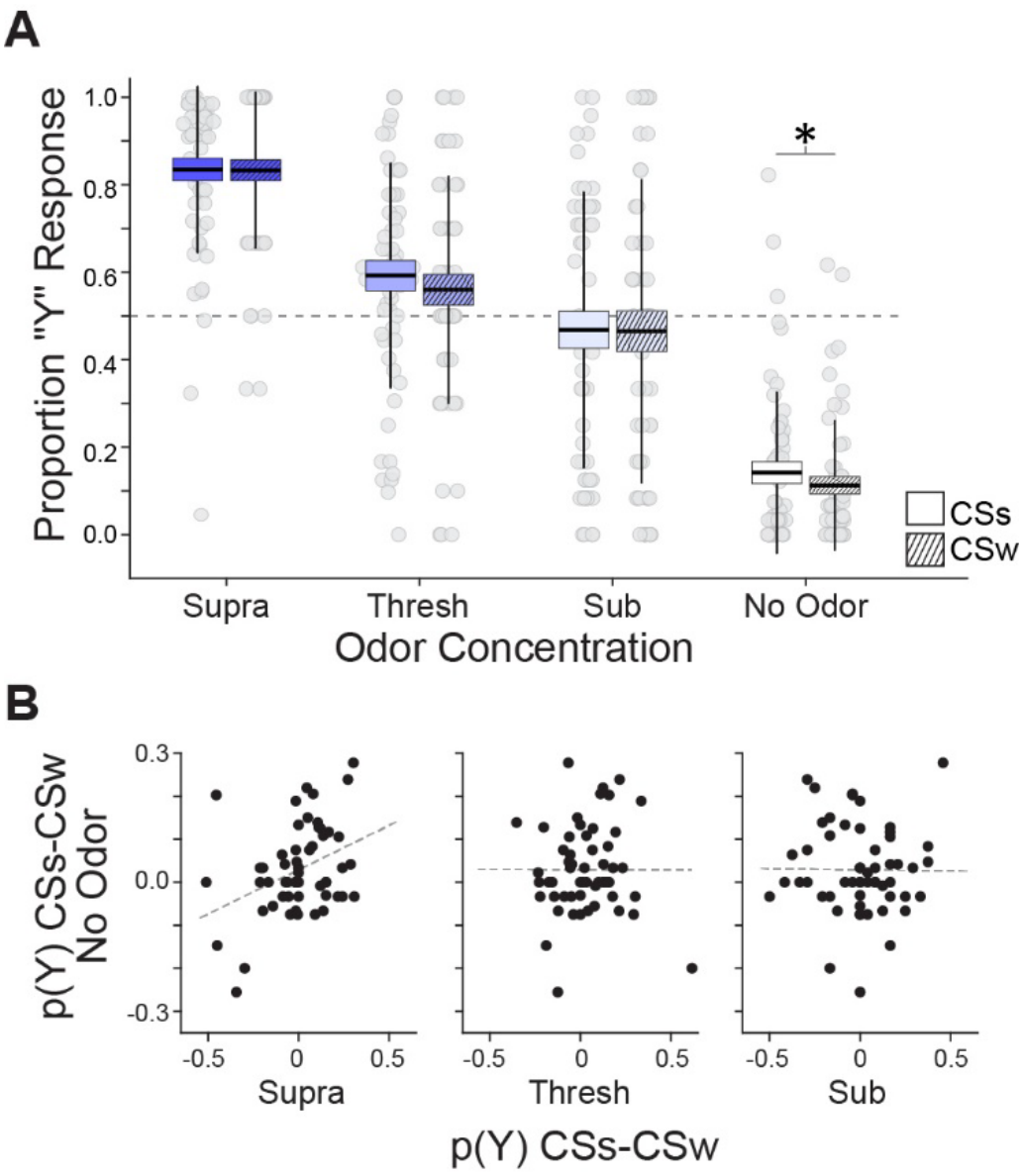
Analysis of odor detection rate. **(A)** The proportion of “Yes” responses (i.e., odor detection rate), was significantly greater on CSs trials than CSw trials in the “no odor” condition, but not different in the supra-threshold, threshold, or sub-threshold conditions. Horizontal lines depict the mean across participants, vertical lines depict standard deviation, the extent of the boxes depict standard error of the mean, and light gray circles depict individual participants. **(B)** The difference between CSs and CSw odor detection rate in the “no odor” condition was correlated with difference in detection rate in the supra-threshold condition (left scatter plot), but not threshold concentration (middle), or sub-threshold concentration (right). p(Y) = proportion “Yes” response. Black circles depict individual participants, and dashed lines depict the linear trend line.

In terms of confidence ratings, we first compared ratings for “Yes” and “No” responses within each concentration step, averaged across cue type. Confidence was significantly higher for “Yes” responses than “No” responses in the supra-threshold condition (t_48_ = 2.94, *p* = 0.0049), and lower for “Yes” than “No” responses in threshold (t_51_ = −3.33, *p* = 0.0016), sub-threshold (t_50_ = −5.59, *p* = 9.30×10^−7^), and “no odor” conditions (t_43_ = −9.35, *p* = 6.45×10^−12^)(**Fig. 4A**). These results demonstrate that, in general, participants used the confidence scale to indicate uncertainty of odor presence in accordance with the concentration of the odor, such that higher concentrations elicited more confidence “Yes” responses and lower concentrations elicited more confident “No” responses. Within the “no odor” condition specifically, participants made more confident “Yes” responses on CSs trials than CSw trials (t_30_ = 2.19, *p* = 0.037) but were equally confident in their “No” responses (t_54_ = 0.74, *p* = 0.46) (**Fig. 4B**).

**Figure 4.**
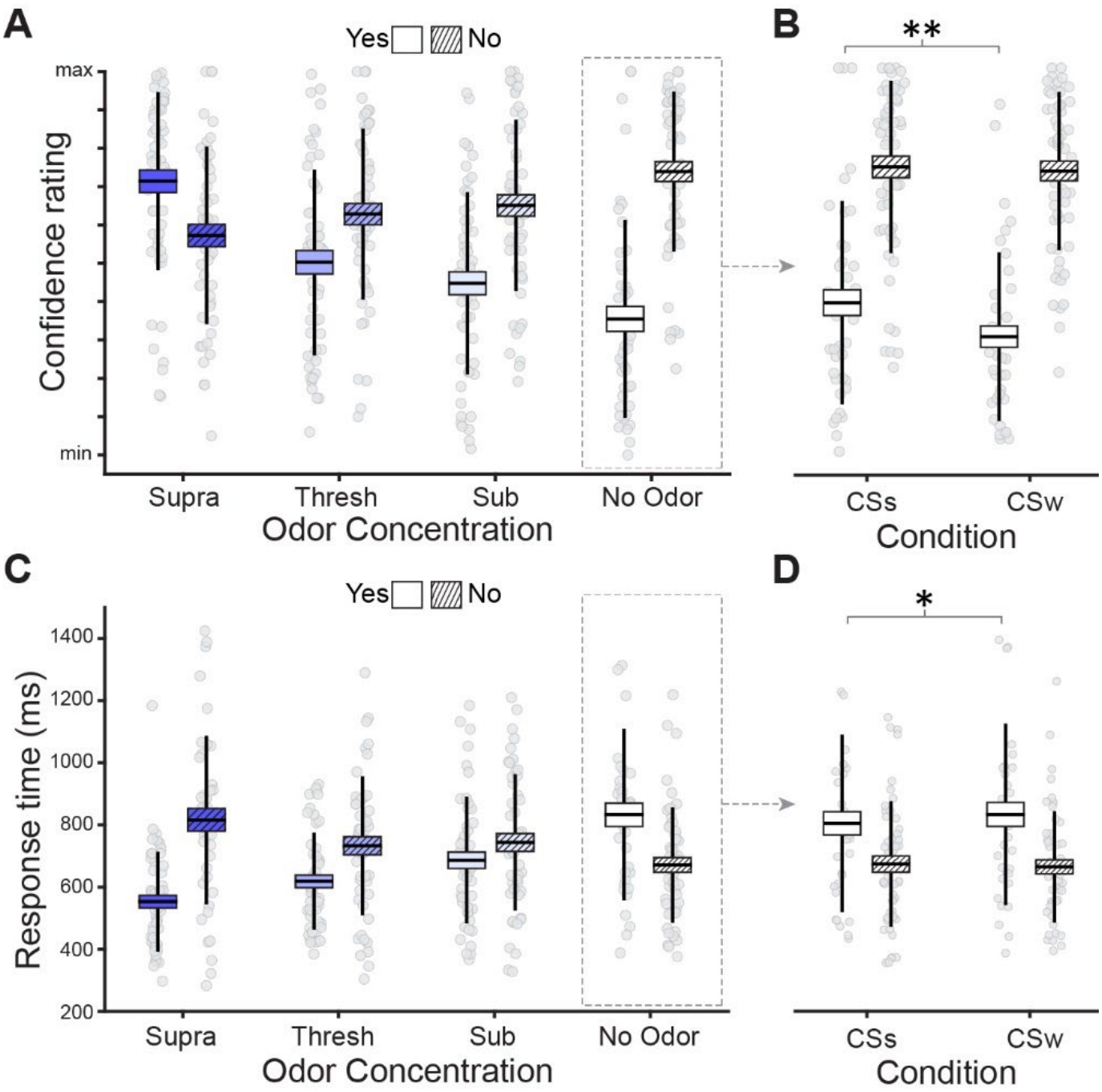
Odor detection task confidence ratings and response times. **(A)** Confidence is plotted by odor concentration and response type, averaged across cue types and all trial blocks. **(B)** Confidence is plotted by response type and CS type specifically in the “no odor” condition. **(C)** Response time is plotted by odor concentration and response type, averaged across cue types and all trial blocks. **(D)** Response time is plotted by response type and CS type specifically in the “no odor” condition. In all panels horizontal lines depict the mean across participants, vertical lines depict standard deviation, the extent of the boxes depict standard error of the mean, and light gray circles depict individual participants.

Averaged across cue types, response times were significantly faster for “Yes” responses than “No” responses in the supra-threshold (t_43_ = −6.34, *p* = 1.15×10^−7^) and threshold (t_46_ = −3.47, *p* = 0.0011) conditions, and marginally faster in the sub-threshold condition (t_47_ = −1.82, *p* = 0.075). In the “no odor” condition “No” responses were significantly faster than “Yes” responses (t_38_ = 3.09, *p* = 0.0038) (**Fig. 4C**).Thus, similar to confidence ratings, participants showed a general trend for faster “Yes” responses at higher odor concentrations, and faster “No” responses at lower concentrations. Within the “no odor” condition specifically, participants made marginally faster “Yes” responses on CSs trials than CSw trials (t_27_ = 1.78, *p* = 0.087), and there was no difference in response time for “No” responses (t_49_ = −0.52, *p* = 0.60) (**Fig. 4D**). Taken together with the detection rate and confidence rating results, these findings demonstrate that participants made more frequent, more confident, and marginally faster false odor detections in the CSs condition compared to the CSw condition.

At the end of the experiment participants rated how certain they were that each CS was paired with a detectable odor during the experiment, on a 5-point scale ranging from “sure no odor” to “sure odor.” Participants rated the CSs as having a higher certainty of odor presence (*M* ± *SD* = 3.35 ± 0.99) than the CSw (*M* ± *SD* = 2.76 ± 0.98; t_54_ = −2.93, p = 0.005) (**Fig. 5A**). However, there was no correlation between the difference in odor certainty rating between CSs and CSw and the difference in response rates for these cues in the “no odor” condition (*r* = −0.0183, *p* = 0.89, Spearman rank) (**Fig. 5B**). Moreover, there was no correlation between the difference and response rate between CSs and CSw in the “no odor” condition and any of the self-report questionnaires measures we acquired (see Methods, *p*’s > 0.36). The observed effect of greater false odor detection rate for the CSs condition is thus unlikely to be driven by a conscious awareness of the increased pairing of that cue with higher odor concentrations compared to CSw, or to individual differences in mental imagery or other potentially clinically relevant factors such as propensity to experience anxiety, depression, hallucinations, or delusions.

**Figure 5.**
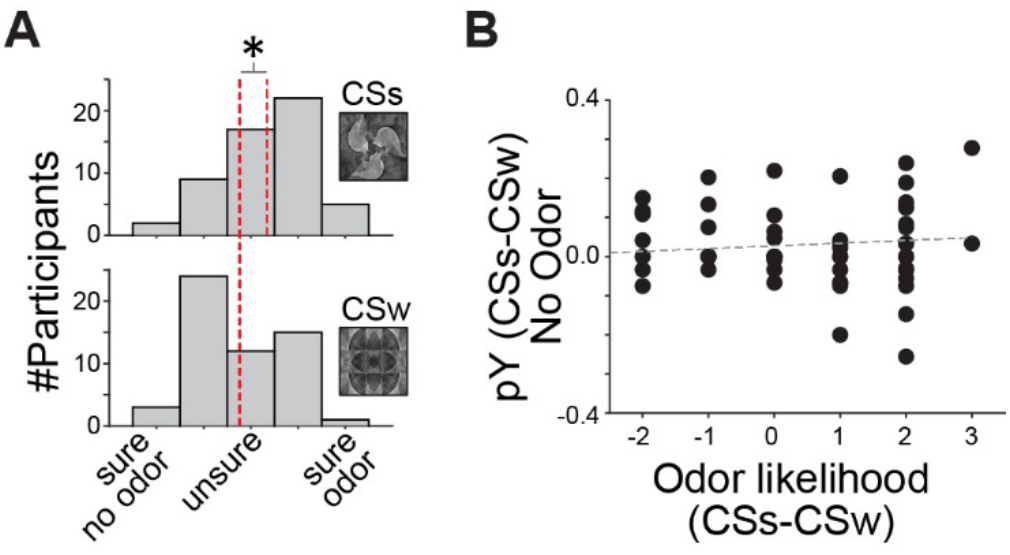
Odor certainty ratings. **(A)** Histograms show certainty ratings for the CSs and CSw conditions for all participants. Red dashed lines indicate the mean rating across participants. **(B)** Scatter plot shows correlation between certainty ratings and false odor detection rate. Circles are individual participants, and the dashed line depicts the linear trend.

## Discussion

Our novel odor detection task demonstrates that robust learned associations between previously neutral stimuli can bias perception such that the presentation of a predictive visual cue alone can result in the false perception of its associated odor. While conditioned hallucination paradigms have demonstrated similar effects in other sensory modalities (Kafadar et al., 2022; Koh & Gallagher, 2020; Powers et al., 2017), this is the first study to replicate this phenomenon in the olfactory domain. Further, by adapting the conditioned hallucination paradigm to include two conditioned stimuli with different associative strengths, we directly examined the influence of learned associations on perception. Following repeated pairings between the strongly conditioned stimulus (CSs) and supra-threshold and threshold odor concentrations, participants were significantly more likely to report detecting an odor during ‘no-odor” trials compared with the weakly conditioned stimulus (CSw). Interestingly, participants expressed greater confidence in these false detections for the CSs, suggesting to some extent a belief that an odor was present. Although a post-task odor likelihood rating did not directly correlate with responses during the “no odor” condition, participants nevertheless subjectively reported believing that the CSs was more likely to be paired with a detectable odor compared with the CSw. This finding suggests that the association established during the initial conditioning phase persisted throughout the latter half of the experiment, even after both conditioned stimuli were paired equally with the no odor concentration. Collectively, these findings indicate that associative learning influenced the participants’ perceptual experience.

These findings are consistent with a predictive processing account of perception, which increasingly conceptualizes perception as an inferential process in which prior expectations are integrated with incoming sensory information to generate perceptual experiences (Teufel et al., 2015; Wilkinson, 2014; Zarkali et al., 2019). Olfaction may be particularly susceptible to this computational framework as odor stimuli are often ambiguous, allowing for multiple possible causes to account for the same sensory input (Grabska-Barwińska et al., 2017). This uncertainty increases reliance on prior expectations during perceptual inference and may render the olfactory system especially vulnerable to prediction errors when learned expectations conflict with incoming sensory evidence. Under these conditions, misattributed sensory evidence may become incorporated into new learning, ultimately giving rise to false perceptions. The present findings support the notion that robust cue-odor associations can bias perceptual inference such that internally generated expectations are experienced as genuine sensory events.

The present findings also contribute to broader theories of hallucinations. Hallucinations occur across all sensory modalities and often take place in the absence of pathology (Allen et al., 2005; Johns et al., 2004; Rogers et al., 2021). Nevertheless, the majority of hallucination research has centered on clinical auditory and visual hallucinations, as these represent the most common hallucination types (Lim et al., 2016; McCarthy-Jones et al., 2017). Comparatively, olfactory hallucinations (OHs) remain much less studied (Fry et al., 2022) despite OHs spanning the broad spectrum of odors encountered in everyday life and being documented in both healthy and various clinical populations (Arguedas et al., 2012; Bainbridge et al., 2018; Ohayon, 2000; Wehling et al., 2021).

Predictive processing theories propose that hallucinations arise when perceptual priors become overweighted relative to incoming sensory evidence. Research further suggests that this overweighing is not characteristic of psychotic disorders specifically but appears to be representative of individuals who experience hallucinations more broadly (Corlett et al., 2019; Powers et al., 2017). Moreover, individuals who do not ordinarily hallucinate can nevertheless experience hallucinations in an experimental setting (Ellson, 1941; Leuba & Dunlap, 1951), supporting the view that hallucinations are an extension of normal perceptual inference rather than a process unique to psychiatric illness (Corlett et al., 2019). It is therefore reasonable to propose that prediction errors, generated by discrepancies between expectations and incoming sensory evidence input, create a need to reconcile these conflicting sources of information. Under conditions in which sensory evidence is weak or ambiguous, strongly weighted expectations may dominate perceptual inference, resulting in internally generated predictions being experienced as genuine percepts. The present findings provide behavioral support for this theoretical framework within the olfactory domain by demonstrating that learned expectations alone are sufficient to bias odor perception in healthy individuals.

Although the present study demonstrates that learned expectations can induce false odor perception, the underlying neural mechanisms remain unknown. Future studies incorporating neuroimaging methodologies could help determine whether participants are engaging neural pathways typically associated with genuine odor perception when reporting false odor experiences. Such work could address not only whether experimentally induced false odor perception engages the olfactory system in a similar fashion to veridical odor perception, but also how prediction errors are resolved during olfactory perceptual inference. These studies would help distinguish the behavioral phenomenon observed here from response-related processes and provide mechanistic insight into how learned expectations influence olfactory perception.

Several limitations should be considered when interpreting these findings. First, the use of a fixed range of odor dilutions resulted in estimated detection thresholds that exceeded the available concentration range for 28 of the 55 participants, with 18 participants falling below the lowest concentration step and 10 above the highest concentration step. The slightly elevated average detection rates in the odor detection task (supra-threshold = 83.4%, threshold = 57.6%, and sub-threshold = 46.7%) likely reflect the overrepresentation of participants whose true threshold fell below the lowest available concentration. Despite this, response patterns closely approximated the expected pattern of performance, suggesting that the odor conditioning remained effective. Second, because odor dilutions were prepared throughout data collection minor variability in concentration across participants cannot be completed excluded. However, rigorous preparation procedures, including precise measurement techniques and standardized experimental methodology minimized this potential source of variability. Finally, a small number of participants (*n* = 5) were included from an earlier iteration of the study. These participants completed the same conditioning paradigm, viewed the same visual stimuli, received the same odor, and completed identical behavioral responses. They differed only in minor refinements to instructions, stimulus timing and odor concentration steps, which is unlikely to have influenced the primary outcome.

Taken together, this study demonstrates that learned associations are sufficient to induce false odor perception in healthy individuals. These findings further provide behavioral support for a predictive processing account of perception by demonstrating how strongly weighted expectations can override incoming sensory evidence to generate genuine false percepts. Clarifying the mechanisms through which perception can go awry has important implications for understanding hallucinations across both healthy and clinical populations. Future studies should replicate these findings using neuroimaging techniques to identify the neural substrates underlying experimentally induced false odor perception to determine how predictive processing within the olfactory system contributes to hallucination-like experiences. Ultimately this could improve our understanding of psychiatric and neurological disorders characterized by false perception while providing broader insight into how expectations shape normal human perception.

